# Lipid Tail Length Determines Nano-Bio Interactions of Peptide Amphiphile Nanostructures

**DOI:** 10.1101/2025.11.18.689169

**Authors:** Karla Bonic, Morgan R. Stewart, Li Xiang, Kailin Mooney, Randall Armstrong, Sadik Esener, Jared M. Fischer, Adem Yildirim

## Abstract

Understanding the interactions of nanomaterials with biological systems is essential to designing effective nanomedicines. However, most of our understanding originates from studies with solid nanoparticles, and nano-bio interactions of self-assembled nanomaterials have remained largely unexplored. To address this knowledge gap, we develop a series of self-assembling peptide amphiphiles (PAs) with different lipid modifications and investigate their interactions with biological systems. We find that PA nanostructures rapidly disassemble and reassemble with albumin and lipoproteins in blood plasma. While PAs with shorter lipid tails mainly assemble with albumin, increasing lipid length shifts binding to lipoproteins. All PAs show strong tumor accumulation in 4T1 tumor-bearing mice with tumor to liver ratios of ∼3-6. Overall, albumin-binding improves blood circulation and tumor accumulation compared to lipoprotein-binding, but also increases off-target accumulation. This study shows that the biointeractions of self-assembled nanomaterials can be controlled through molecular design, which may lead to the development of effective nanomedicines.

## 1. Introduction

Nanoscale carriers, such as liposomes, micelles, or lipid, polymer, or inorganic nanoparticles, are promising for the targeted delivery of contrast agents and drugs to solid tumors as they offer improved cargo solubility and loading, surface functionalization for targeting, and controlled release in the tumor microenvironment.^[1-3]^ Yet only a handful of cancer nanomedicines are successfully implemented in the clinic, the majority being liposomal drug formulations.^[4-6]^ Major challenges against the clinical translation of nanomedicine include a limited understanding of the interactions of nanocarriers with biological systems, control of payload leakage and rapid nanocarrier clearance in circulation, off-target accumulation, and complex and multi-step fabrication.^[7-9]^ A promising alternative of synthetic nanocarriers is to use biological macromolecules such as albumin, lipoproteins, exosomes, or cell membranes.^[10-13]^ These intrinsically biocompatible biological nanomedicines can naturally accumulate in solid tumors as a result of the increased metabolism of cancer cells and their role in nutrient transport and cellular communication.^[14-16]^ While these platforms show promise in preclinical studies, their clinical translation has remained a challenge due to complex biomolecule isolation, drug loading and purification steps, and difficulties in their large-scale and reproducible production.^[17-19]^ An alternative strategy to address these limitations is developing payload-conjugated small molecules that can chemically or physically assemble with biomolecules *in vivo* upon systemic delivery and exploit them to specifically accumulate in solid tumors.^[20,21]^ These small molecule ‘hitchhikers’ can be easily produced at a large scale; thus, they can facilitate clinical translation.^[22-23]^ In fact, an albumin-binding prodrug of doxorubicin (aldoxorubicin) showed promising antitumor activity in clinical trials.^[24]^ Despite this strong premise, the research on hitchhiker development has been limited and remained mainly focused on albumin.^[25]^

In a recent study, we showed that weakly assembled nanostructures of peptide amphiphiles (PAs) could disassemble in circulation and reassemble with blood biomolecules, mainly lipoproteins.^[26]^ Assembly with lipoproteins significantly prolonged blood circulation and enabled strong accumulation and retention in a broad range of solid tumors due to increased lipid metabolism of cancer cells.^[27-30]^ Building upon these initial findings, in this work, we investigated how the PA structure affects their interactions with biological systems **(Figure 1)**. We prepared a series of PAs with different saturated lipid modifications with different tail lengths while keeping the peptide backbone the same to change the overall hydrophobicity of the PAs. We found that while more hydrophilic PAs with short lipid tails mainly assembled with albumin, increasing the hydrophobicity reduced albumin binding and improved assembly with lipoproteins. In addition, increasing the hydrophobicity of the PAs improved their cellular internalization through stronger assembly with cell membranes. Biodistribution studies in orthotopic 4T1 mouse breast tumor-bearing mice found that the interactions with plasma components determine the biodistribution and tumor accumulation of PAs. While binding to lipoproteins and albumin both significantly improved blood circulation, a slower blood clearance was observed for albumin-binding PAs. We also found that blood circulation of PAs is the main determinant for total tumor accumulation. Finally, while albumin-binding PAs showed significantly higher tumor accumulation, albumin-binding is also found to increase accumulation in the liver and other organs compared to lipoprotein-binding PAs. Overall, this work demonstrates that the previously overlooked interactions of self-assembled PA nanostructures with biomolecules determine their biodistribution, and it describes how these interactions can be tuned to control their biodistribution and tumor accumulation.

**Figure 1.**
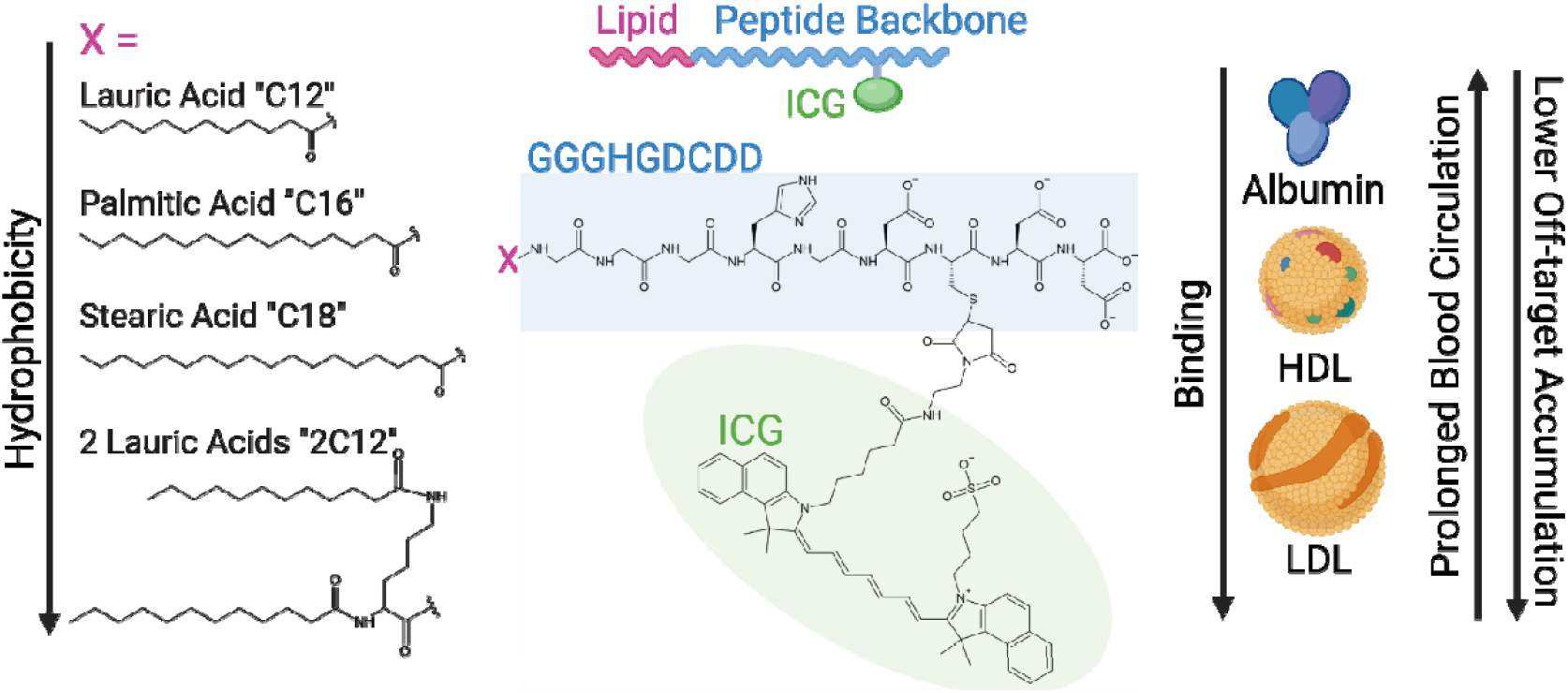
Schematic summarizing the structure of PAs, and proposed effect on biomolecule binding and biodistribution *in vivo*. Created with BioRender and ChemSketch.

## 2. Results and Discussion

To study the effect of lipophilicity of peptide amphiphiles (PAs) on their interactions with endogenous biomolecules, we prepared PAs with different n-terminal fatty acid modifications with different lengths (Figure 1): lauric (C12), palmitic (C16), or stearic acid (C18). In addition, a PA with two lauric acid modifications (2C12) was prepared by adding a lysine to the n-terminal of the peptide sequence. All PAs share the same hydrophilic amino acid sequence (GGGHGDCDD), which is designed based on our previous results.^[26]^ PAs were conjugated with indocyanine green (ICG) dye through thiol-maleimide coupling to track them *in vitro* and *in vivo*. PA-ICG conjugates were purified via high-performance liquid chromatography (HPLC) and characterized using liquid chromatography-mass spectrometry (LC-MS) **(Supporting Information, Figure S1-4)**. The morphology of the self-assembled PA nanostructures in water was investigated using transmission electron microscopy (TEM). All PAs formed self-assembled structures regardless of their lipid modification due to the amphiphilic peptide-based structure, driven by hydrophobic interactions between alkyl chains and hydrogen bond interactions with the peptide terminal **(Supporting Information, Figure S5)**. All PAs formed spherical micelle structures with sizes around 10 nm, which is in accordance with our previous study.^[26]^ More organized and less aggregated structures were observed when the lipid chain length was increased from 12 to 16, most likely due to improved hydrophobic intermolecular interactions in these structures.^[31,32]^ Further increasing the hydrophobicity did not significantly affect the micelle morphology.

First, we investigated how varying lipophilicity affected the structural stability of PAs in human plasma and their interactions with plasma components. To study the stability of PA nanostructures, we incubated PAs or ICG (100 µM) in 10% human plasma and measured fluorescence prior to and at various time points after the addition of plasma. Without plasma, ICG fluorescence was completely quenched for all PAs due to the dense packing of ICG molecules in their self-assembled structures (**Figure 2a**). Upon the addition of plasma, PAs and ICG exhibited an immediate increase in fluorescence, indicating the disassembly of the PA nanostructures. This result was in accordance with our previous report, where a structurally similar negatively charged PA showed complete disassembly in plasma.^[26]^ We also observed that the fluorescence of PAs decreased over time, suggesting another conformational change for PAs, which might happen upon binding to plasma components. In addition, secondary fluorescence quenching became more pronounced with increasing hydrophobicity, indicating stronger intermolecular interactions for these more hydrophobic PAs. Overall, these results showed that all of the PAs developed in this study can effectively disassemble in plasma, but their interactions with plasma components are likely to be different.

**Figure 2.**
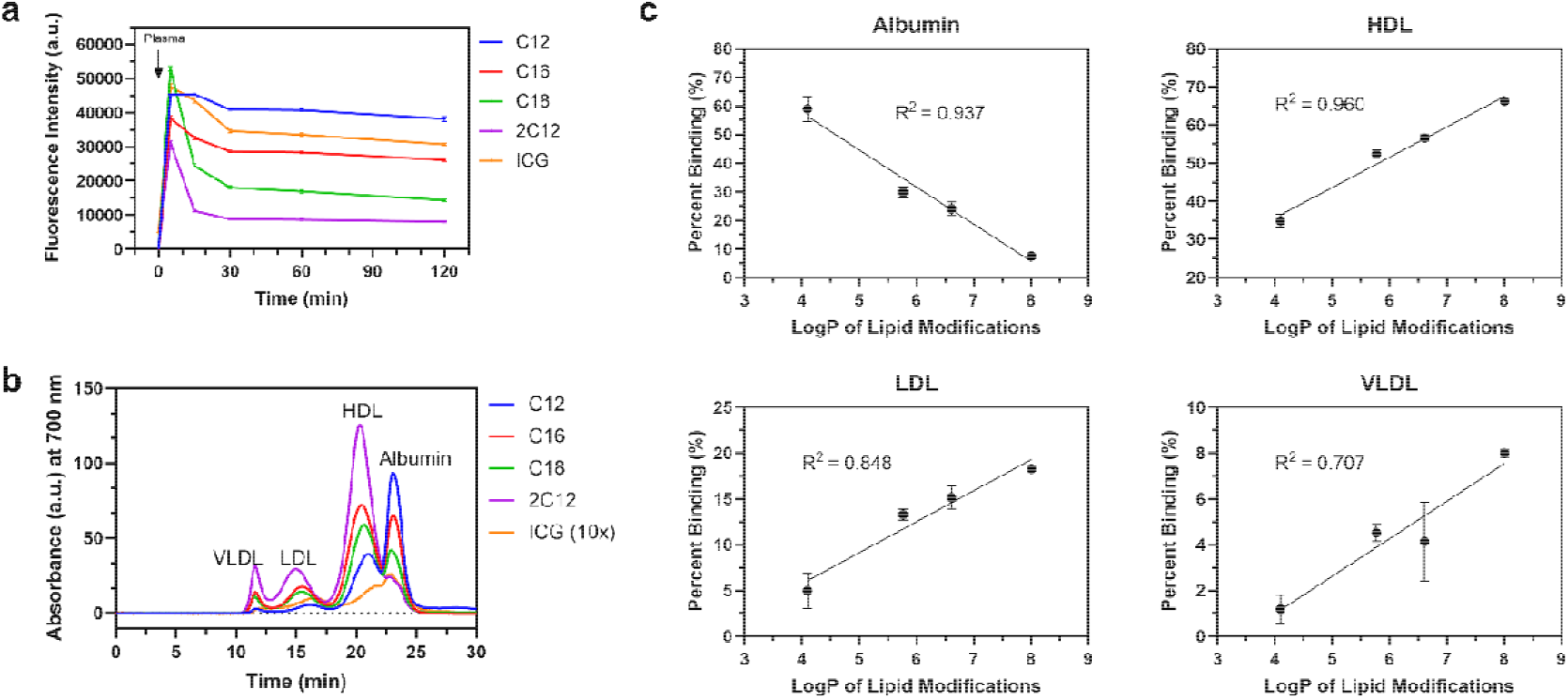
Lipid modification of PAs determines their interactions with blood plasma components. a) Fluorescence intensity of 100 µM PAs prior to plasma addition (plotted at 0 min), and after adding 10% human plasma over two hours. b) FPLC traces of PAs (100 µM) in 10% human plasma detected using absorbance of ICG at 700 nm. To display ICG binding to plasma components more clearly, its absorbance was rescaled by multiplying by 10 (10x). c) Plots showing percent binding of PAs to plasma components, calculated from FPLC traces, versus LogP values of PA lipid modifications. Studies were run in quadruplicate in (a) and triplicate in (c). Data in (a, c) are presented as mean ± standard error of the mean (SEM).

The most likely interactions of PAs are expected to be with albumin and lipoproteins as they are highly abundant in the blood plasma and carry hydrophobic molecules.^[33,34]^ To investigate these interactions, we utilized fast protein liquid chromatography (FPLC) measurements as described in our previous study.^[26]^ We performed FPLC of ICG and PAs in PBS or human plasma and utilized absorbance of ICG at 700 nm to separate PAs from plasma components. Without plasma, micelles of C12 were undetectable **(Supporting Information, Figure S6)**, likely due to the formation larger aggregates that are unable to pass through the FPLC column. More hydrophobic variants eluted in a single peak without plasma suggesting smaller self-assembled structures for these PAs as observed by TEM. When incubated with plasma, C12 eluted mostly with albumin (59.0% ± 4.3%), with some HDL binding (34.9% ± 1.7%) **(Figure 2b)**. A gradual decrease in the albumin binding and an increase in lipoprotein binding was observed with increasing PA lipophilicity. We also calculated the octanol-water partition coefficient (LogP) values of lipid modifications using ChemDraw software (v23.1.1, Revvity), which showed a good linear relationship between LogP of lipid modifications and percent binding for all lipoproteins and albumin **(Figure 2c)**. In accordance with our previous work, C16 eluted mostly with HDL (52.4% ± 1.0%) and, to a lesser extent, with albumin (29.8% ± 1.9%). Also, binding to low-density lipoproteins became significant, with C16 eluting with LDL (13.3% ± 0.6%) and VLDL (4.5% ± 0.4%). For C18 and 2C12, lipoprotein binding became more pronounced, averaging 75.9% and 95.5%, respectively. In addition, overall area under the curve (AUC) values of FPLC plots increased gradually with increasing PA lipophilicity (198.0 ± 27.6, 368.1 ± 78.7, 242.4 ± 10.8, 453.1 ± 8.2, for C12, C16, C18, and 2C12, respectively), except for C18, suggesting that PAs with more hydrophobic lipid modifications can bind more strongly to the plasma components. The lower AUC observed for C18 compared to C16 suggests that the self-assembled structures of C18 remained partially stable. Finally, ICG eluted at a very slight peak with albumin. However, the overall AUC was 18.5-fold lower than that of C12, indicating that only a small percentage of ICG assembles with plasma components. Overall, these results showed that the plasma components that PAs assemble with can be finely tuned by changing their lipophilicity and it can be estimated from LogP value of lipid modification.

After showing that lipid tail length of PAs determines their interactions with plasma components, we wanted to explore how structural differences of PAs affect their interactions with cell membranes and cellular internalization. For these studies, 4T1 mouse breast cancer cells were treated with 10 µM ICG labeled PAs for two hours and imaged using a confocal microscope **(Figure 3a)**. While all PAs were taken up by cells, significant retention in the membrane was observed for PAs with longer lipid tails (C16, C18, and 2C12), suggesting that PAs internalize in cells through membrane binding of their monomers, not as intact micelles. In addition, we observed that the internalization of PAs increased with increasing lipophilicity except for 2C12 **(Figure 3b)**. To gain more insights on the cellular uptake of PAs, we treated 4T1 cells for 1 min with 50 µM of PAs and then imaged the cells at different time points **(Supporting Information, Figure S7)**. All PAs initially showed membrane binding and then internalized in cells over time. We observed that more hydrophobic PAs stayed on the cell membrane longer. We also explored if strong membrane binding of PAs causes any toxicity and found no significant decrease in the viability of 4T1 cells incubated with the PAs for three days, except for 2C12 at concentrations higher than 10 µM **(Supporting Information, Figure S8)**. The slight toxicity observed for 2C12 can be due to the disruption of the cellular membrane at high PA concentrations. Altogether, these studies suggest that the PAs internalize in cells without causing significant toxicity.

**Figure 3.**
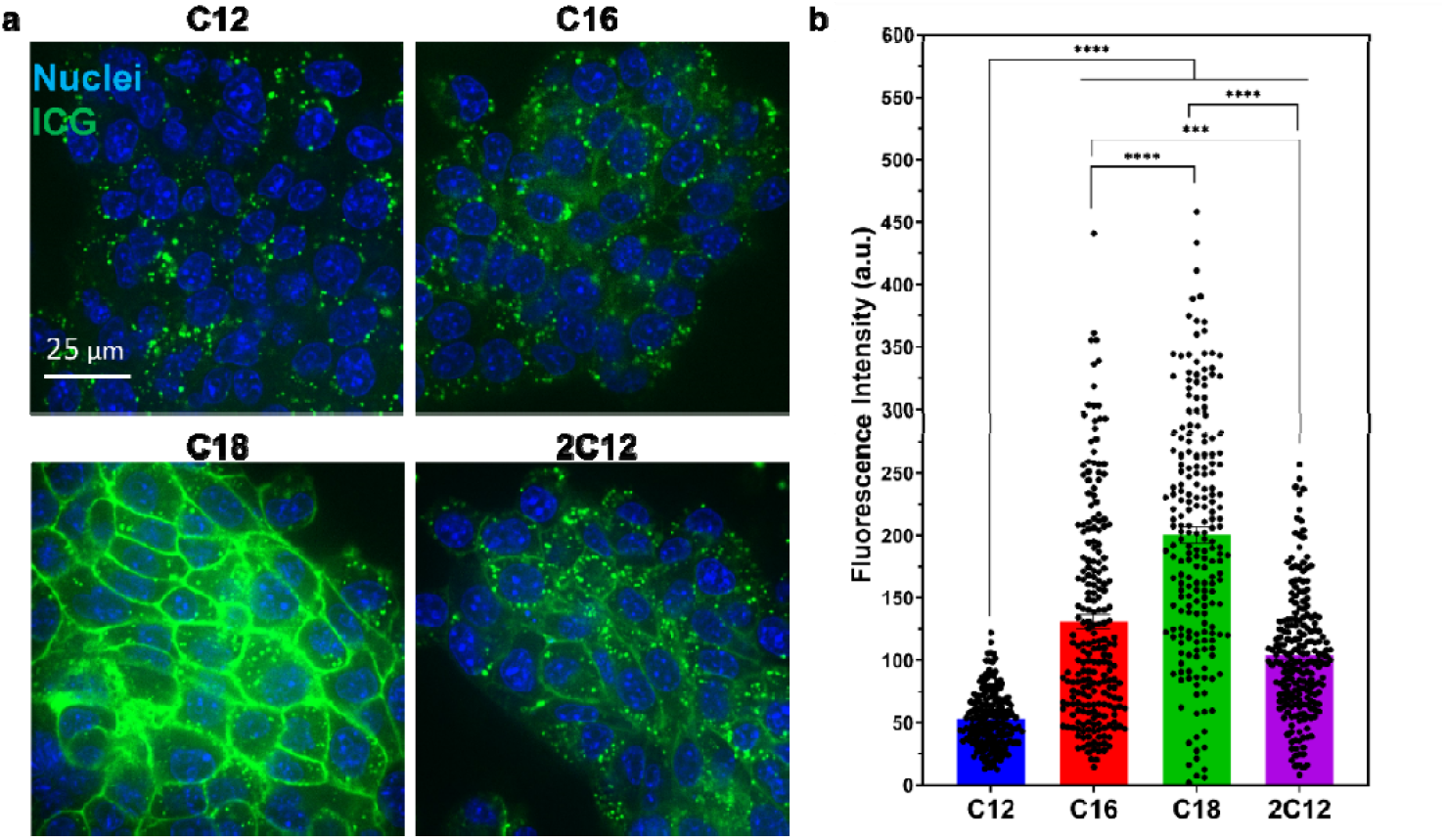
PAs assemble with cell membranes for cellular internalization. a) Confocal microscope images of 4T1 cells treated with PAs (10 µM) for 2 hours. Green = ICG, Blue = Nuclei. b) Quantitative uptake of PAs in 4T1s. All studies were run in quadruplicate. Data are presented as mean ± standard error of the mean (SEM). Statistical analysis was performed using one-way analysis of variance (ANOVA), \*\*\**p* <0.001, \*\*\*\**p* <0.0001.

Having found significant differences in the interactions of PAs with blood plasma biomolecules and cell membranes, we moved to *in vivo* studies to explore how these interactions affect the biodistribution and pharmacokinetics of PAs. For these studies, we intravenously injected PAs (50 nmole) into mice bearing 4T1 mCherry mammary tumors. We collected blood samples from the mice up to 2 days to study their blood circulation. Mice were also imaged using IVIS for up to 2 days to determine tumor accumulation and background signal. At the end of the experiment, the organs of mice were harvested and imaged using IVIS to determine the biodistribution of the PAs **(Figure 4a)**. All PAs showed largely prolonged blood circulation compared to free ICG injection, with blood circulation half-lives of 4.9, 3.9, 3.1, and 3.1 hrs for C12, C16, C18, 2C12, respectively, compared to <5 min for ICG ^[35]^ **(Figure 4b)**, showing that assembly with plasma components improves blood circulation. In addition, we found that C12, which mainly binds to albumin, showed significantly slower blood clearance compared to lipoprotein binding PAs (C16, C18, and 2C12), which can be explained by longer blood circulation half-life of albumin (∼20 days) than lipoproteins (∼0.5-3 days).[36-38]

**Figure 4.**
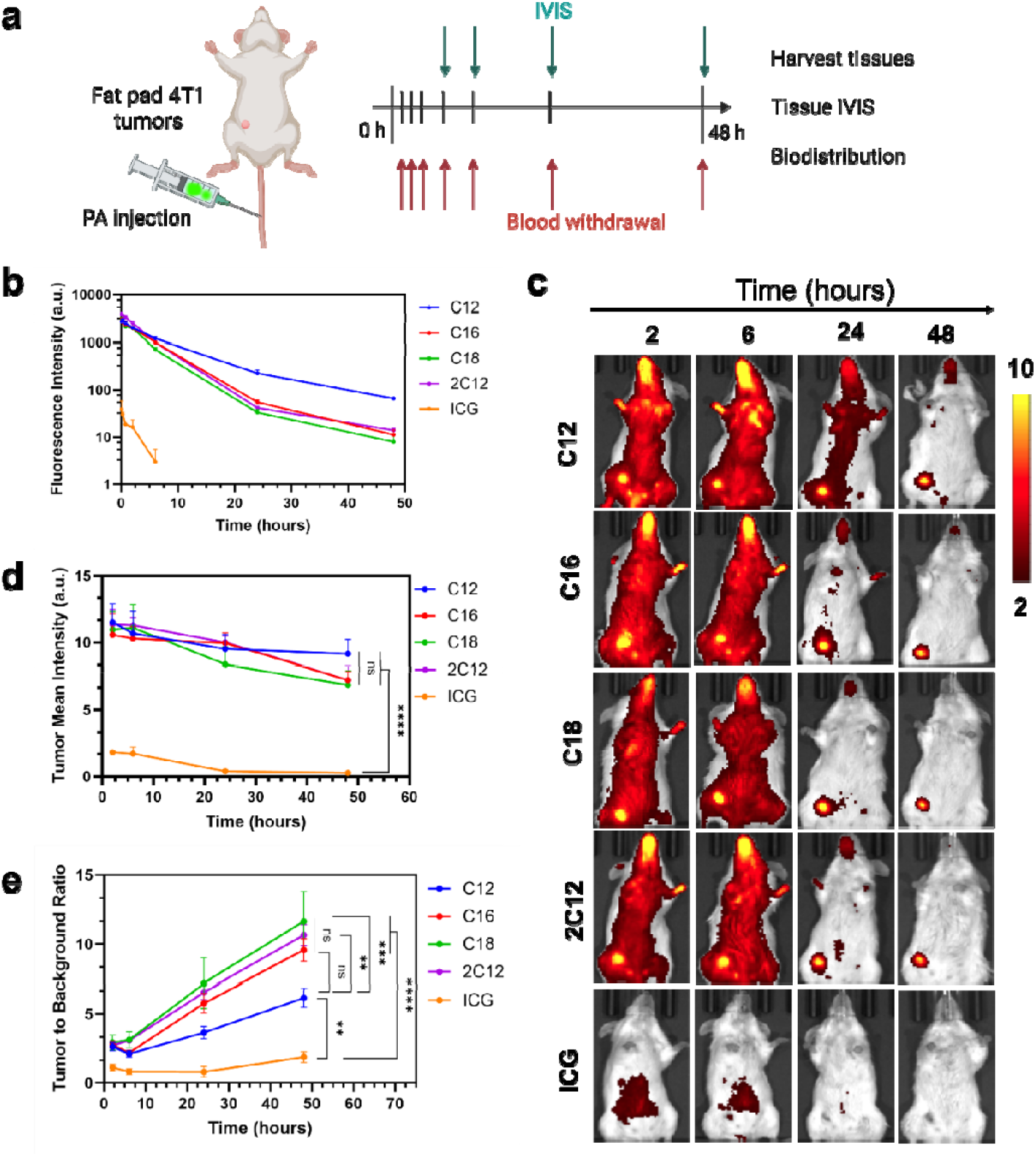
Assembly of PAs with plasma biomolecules enables prolonged blood circulation and enhanced tumor accumulation compared to free ICG. a) Schematic showing the experimental methods used to determine the effect of PAs on tumor specific delivery. Created with Biorender.com. b) Blood circulation of PAs or free ICG (50 nmole) after tail-vein injection in 4T1 tumor bearing mice. c) Representative IVIS images and d) ICG signal in the tumor and e) tumor-to-background ratio after tail-vein injection of PAs or ICG (50 nmole) in 4T1 orthotopic mice at different time points show significantly higher tumor and tumor-to-background signal for all PAs versus ICG. All studies were run in quadruplicate except for ICG-injected mice which were done in triplicate. Error bars are standard error of the mean (SEM). Statistical analysis was performed using two-way analysis of variance (ANOVA). *n*.*s*. is non-significant and \**p* <0.05, \*\**p* <0.01, \*\*\**p* <0.001, \*\*\*\**p* <0.0001.

Next, we studied the tumor accumulation of the PAs over time. IVIS on live animals showed similar tumor accumulation kinetics for all PAs with strong tumor accumulation in the first 2 hours **(Figure 4c, d)**. The tumor signal remained strong over 2 days with a slight decrease of around 10-30% compared to the intensity 2 hours post-injection for all PAs. In accordance with our previous results, free ICG injection did not highlight the tumor at any of the time points.^[26]^ While a strong tumor signal was observed 2 hours after injection, the tumor-to-background ratio (TBR) was still >2 for all PAs **(Figure 4e)**. The background signal was cleared over time, and TBR reached >10 for lipoprotein binding PAs (C16, C18, and 2C12). Background signal of albumin binding C12 was significantly higher than other PAs on day 2, which resulted in a lower TBR of 6. For free ICG, TBR remained almost constant at around 1 throughout the experiment. These results suggested that PAs that mainly bind to lipoproteins can detect solid tumors more specifically in live animals compared to PAs that primarily assemble with albumin.

Next, we investigated the biodistribution of PAs using IVIS on the tissues collected 2 days post PA injection **(Figure 5a and Supporting Information, Figure S9)**. Consistent with the live animal imaging, all PAs showed similar tumor accumulation and albumin binding C12 showed higher signals in other organs, especially in the liver, kidney, and lung, compared to other PAs. To investigate the accumulation of PAs at the cellular level in the tumor microenvironment, tumor sections were imaged using confocal imaging **(Supporting Information, Figure S10)**. All PAs showed mostly uniform distribution in the tumor microenvironment and strong accumulation in mCherry-positive 4T1 cells. Different from IVIS observations, the fluorescence signal of C12 was stronger than other PAs.

**Figure 5.**
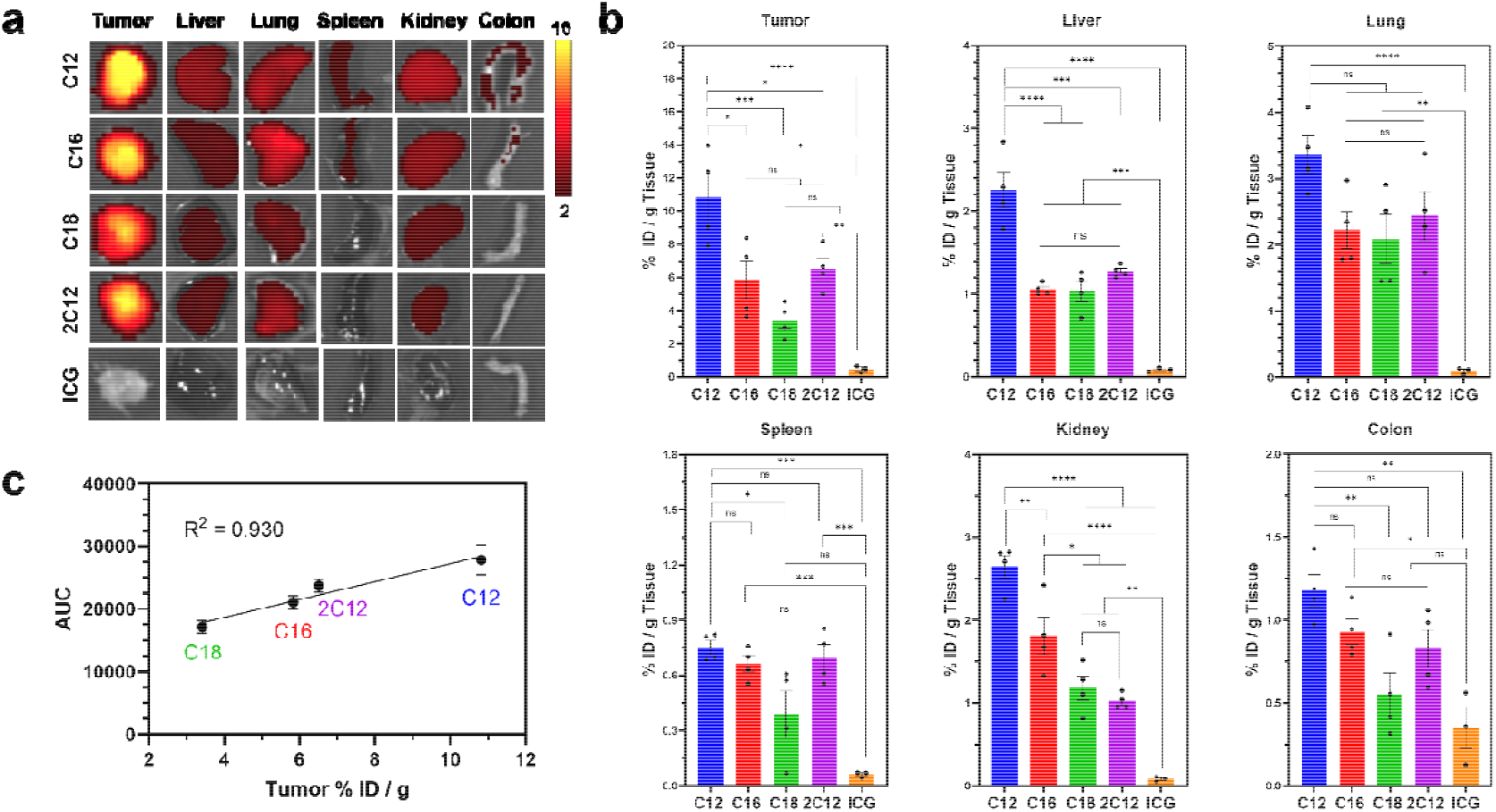
Biodistribution of PAs in 4T1 tumor bearing mice. a) IVIS images of excised tumor, liver, lung, spleen, kidney, and colon two days after intravenous injection of PAs or ICG (50 nmole). b) Percent injected dose / g tissue values of PAs detected in homogenates of tumors and other organs. c) AUC of blood circulation plots of PAs versus percent injected dose found in tumors. Circulation time of PAs is linearly correlated with their relative accumulation in tumors. All studies were run in quadruplicate except for ICG-injected mice which were done in triplicate. Error bars are standard error of the mean (SEM). Statistical analysis was performed using one-way analysis of variance (ANOVA) in (b, c). *n*.*s*. is non-significant and \**p* <0.05, \*\**p* <0.01, \*\*\**p* <0.001, \*\*\*\**p* <0.0001.

To quantify PA accumulation in different tissues, we homogenized tumors and major organs and determined the PA amount in each tissue using calibration curves obtained by spiking C16 to tissue homogenates from a mouse that did not receive any PA injection **(Supporting Information, Figure S11)**. While IVIS of both live and ex vivo showed similar tumor accumulation for all PAs, the ICG fluorescence in homogenized tumor tissues showed significant differences **(Figure 5b)**. Nevertheless, IVIS and tissue homogenization results were comparable in other tissues with significantly higher concentrations for albumin binding C12. The difference between IVIS and fluorescence detection of PAs in homogenized tissues can be explained by the fact that IVIS is a semi-quantitative method due to the differences of light attenuation in different tissues and the signal dependence on tissue thickness (i.e., fluorophores closer to the surface look brighter).^[39]^ In accordance with confocal imaging, a significantly higher tumor concentration was observed for C12 compared to other PAs. In addition, C18 showed significantly lower tumor accumulation than other PAs, which might be due to its shorter blood circulation time (Figure 4b) and lower affinity to plasma components (Figure 2b) with an AUC value 1.5 fold lower than C16. Remarkably, for all PAs, the percent injected dose in gram tissue (%ID / g tissue) of >4% was observed even after 2 days after injection, which reached ∼10% for C12. To understand if there is a correlation between blood circulation time and tumor accumulation of PAs, we calculated the AUC of blood circulation plots of PAs. We then plotted AUC values against tumor % ID / g tissue values, which showed a strong correlation with a linear regression value of 0.930 **(Figure 5c)**. Finally, all PAs showed tumor to liver ratios of around 5-6, except for C18, which was slightly lower, ∼3.3, although not statistically significantly different than others **(Supporting Information, Figure S12)**. Similar trends were also observed for tumor to kidney and lung ratios (Supporting Information, Figure S12). Overall, these results showed that while binding to albumin improved tumor accumulation compared to lipoproteins binding PAs, it also increased the background signal and accumulation in major organs. Nevertheless, all PAs showed favorable biodistribution with strong tumor accumulation and lower accumulation in other tissues.

## 3. Conclusion

In summary, this work showed that self-assembled nanostructures of PAs could quickly disassemble in blood plasma and assemble with major plasma components, albumin and lipoproteins, and hitchhike on them to specifically accumulate in solid tumors. The plasma component that the PAs predominantly assembled with was found to be dependent on the lipid modification of PAs. Overall, more hydrophobic lipid modifications decreased albumin binding and increased lipoprotein binding. This study also showed that more hydrophobic lipid modifications resulted in a stronger assembly with cell membranes, enabling effective cellular internalization *in vitro* and *in vivo*. While both albumin or lipoprotein binding PAs showed strong and selective accumulation in an orthotopic mouse breast cancer model, notable differences in their biodistribution and pharmacokinetics were observed. In general, albumin binding improved blood circulation and tumor accumulation, and assembly with lipoproteins reduced the accumulation in other organs. The findings of this study represent a straightforward yet effective way to control the biodistribution of self-assembled nanomaterials, which may enable specific delivery of medicine to solid tumors to outcomes of cancer patients.

## 4. Materials and Methods

### Materials

Peptides were purchased from Genscript with purity >95%. The following reagents were used: ICG (TCI Chemicals I0535-100MG), ICG maleimide (MedChem Express HY-D1744 or BioActs PWM1301), pooled human plasma (Innovative Research IPLAWBK2E50ML), and anhydrous DMSO (Sigma-Aldrich 276855-100ML).

### Preparation of PA-ICG conjugates

PAs were mixed with 1.1 molar equivalent maleimide-functionalized ICG maleimide in anhydrous DMSO overnight at room temperature with gentle shaking at 400 RPM. The product molecular weight was confirmed by LC-MS using an Acquity UPLC system (Waters) with a SQ Detector 2 (Waters) and a reverse-phase column (Waters, ACQUITY UPLC BEH C18 Column, 130Å, 1.7 µm, 2.1 mm x 50 mm). The LC-MS protocol established a solvent gradient from water/acetonitrile (95:5) with 0.1% formic acid to water/acetonitrile (0:100), run over 9 minutes at a rate of 0.4 mL min^-1^. The product was purified by HPLC (Waters 1525, binary HPLC pump) with a reverse-phase column (Waters, XBridge BEH C18 OBD Prep Column, 130Å, 5 µm, 19 mm x 150 mm). The solvent gradient in the HPLC method consisted of an adjustment from water/acetonitrile (95:5) to (5:95) with 0.1% trifluoroacetic acid over a 45-minute run at a flow rate of 0.8 mL min^-1^. A UV/Visible detector (Waters 2489) measured the absorbance of eluents at 230 nm. Finally, the HPLC purified product was lyophilized, redissolved in filtered PBS and 2.5% DMSO at a concentration of 1 mM and stored at −20 °C.

### TEM imaging of Pas

Formvar/Carbon 300 mesh grids 63 µM holes (Ted Pella 01753-F) were treated using a Pelco EasiGlow 91000 Glow Discharge Cleaning System. PA-ICG conjugates were diluted to 150 µM in ultrapure water. Grids were placed on top of 15 µL drops of the PAs and sat for 10 minutes, then moved to sit in ultrapure water for 3 minutes. 0.4% uranyl acetate (Electron Microscopy Sciences 22400) was added dropwise onto parafilm, and the grids were then placed on top of the drops and let to sit for 5 minutes. Grids were blotted to remove excess stain, and they were left to dry at room temperature. Samples were imaged with TEM using the FEI Tecnai with Icorr.

### PA disassembly study

ICG or PA-ICG conjugates were diluted to 100 µM in filtered PBS, and 90 µL was added to wells of a 96 well plate. Fluorescence spectra of ICG (excitation 720 nm, emission 770-860 nm) were collected using TECAN Spark 20M microplate reader. Then, 10 µL pooled human plasma was added to each well. Fluorescence was measured again at 15, 30, 60, and 120 min post-plasma addition. The plate was incubated at 37 °C 5% CO_2_ between measurements.

### FPLC analysis of PA plasma component interactions

400 µL solutions of ICG or PA-ICG conjugates (100 µM in filtered PBS), with or without 10% human plasma were incubated at 37°C for 30 minutes under gentle shaking at 300 RPM. The samples were then run by FPLC on the GE LJKTA pure™ Chromatography System through a Superose 6 Increase 10/300L column (Cytiva 29091596). Conditions were as follows: 30 mL column volume (CV), 5 MPa pre-column pressure limit, 2.6 MPa delta-column pressure limit, 0.75 mL min^-1^ flow rate, eluted in 100% PBS. The method constituted of an initial equilibration step prior to injection of sample with 100% PBS for 0.5 CV, then sample injection via 1 mL capillary loop in 100% PBS, then finished with a linear elution consisting of 2 CV collected in 1 mL fractions. The UV-Vis detector detected eluted plasma biomolecules at 280 nm and PA-ICG conjugates or ICG at 700 nm.

### Cell culture

4T1 cells (ATCC CRL-2539) were cultured in RPMI 1640 Medium (Gibco 11875093) supplemented with 10% Fetal Bovine Serum (Cytiva, SH30396-03) and 1% 10,000 U mL^-1^ Penicillin-Streptomycin (Gibco 15140163) and incubated at 37°C 5% CO_2_. For mCherry transfection of 4T1 cells, lentivirus targeting mCherry to the cell membrane (Takara, 0026VCT, rLV.EF1.mCherry-Mem-9) was followed from the manufacturer’s protocol. Cells were passaged at least 10 times before using.

### Cellular uptake studies

5-15 ×10^3^ 4T1 cells per well were seeded in a 96 well black plate with glass flat bottom (CellVis P96-1.5H-N) and incubated at 37°C 5% CO_2_ until reaching near confluency. Then, culture media was replaced with 100 µL of 10 µM ICG or PA-ICG conjugates in media and incubated for 2 hours at 37°C 5% CO_2_. All treatment was removed, and cells were washed with PBS, and either incubated with fresh culture media or proceeded to imaging. Fresh media containing 2.5 µg mL^-1^ Hoechst 33342, Trihydrochloride, Trihydrate (Fisher Scientific H3570) was placed on the cells and incubated for 15 minutes prior to imaging. For the time-dependent uptake study, 50 µM ICG or PA-ICG conjugates in media were first incubated at 37°C 5% CO_2_ for 20 minutes. The culture media in the wells was replaced with 100 µL 2.5 µg mL^-1^ Hoechst 33342, Trihydrochloride, Trihydrate and incubated for 15 minutes. Then, 100 µL of the previously incubated solutions replaced the current media and sat for 1 minute. The treatment was then removed and washed with PBS. The wells were then imaged at 1 minute, 5 minutes, 15 minutes, and 90 minutes post-removal of the treatment. Wells were imaged using the Nikon CrestOptics X-Light V3 Spinning Disk Confocal using NIS-Elements software, under a warming chamber also providing 5% CO_2_. Hoechst staining was imaged at 405 nm, 25% laser power, 100 ms exposure. ICG was imaged at 748 nm, 30% laser power, 300 ms exposure. Cellular internalization was quantified by analyzing fluorescence intensity of the PAs with ImageJ.

### PA cytotoxicity

1 ×10^4^ 4T1 cells per well were seeded in a 96-well plate in 100 µL of media. The plates were incubated at 37 °C, 5% CO_2_ for 1 day. Then, media was replaced with 100 µL of PA-ICG conjugates in media at concentrations from 0.1 to 100 µM. Treated plates were incubated at 37 °C, 5% CO_2_ for 3 days. Then, media was replaced with fresh media containing 10% CellTiter 96 Aqueous One Solution Cell Proliferation (MTS) reagent (Promega G3580), and plates were incubated at 37 °C, 5% CO_2_ for 1.5 hours before measuring the absorbance at 490 nm using the TECAN Spark 20M microplate reader.

### Live IVIS study

All experiments were approved by Oregon Health and Science University (OHSU) Institutional Animal Care and Use Committee (IACUC) (TR01_IP00000674) and conformed to the guidelines set by United States Animal Welfare Act and the National Institutes of Health. Mice between 2-4 months and 20-30 g in weight were housed in specific pathogen free cages. 0.5×10^6^ 4T1 mCherry cells were injected into the fat pad of Balb/c mice (The Jackson Laboratory, 000651) and allowed to grow into tumors around 0.5-1 cm. 100 µL of 0.5 mM ICG or PA-ICG conjugates were injected intravenously (IV). Then, mice were imaged using the IVIS Spectrum (PerkinElmer) with excitation and emission wavelengths of 745 and 820 nm, respectively, at the specified time points for up to 7 days. Living Image (PerkinElmer) software was used to analyze fluorescent signal.

### Blood circulation and biodistribution study

0.5×10^6^ 4T1 mCherry cells were injected into the fat pad of Balb/c mice and allowed to grow into tumors around 0.5-1 cm. 100 µL of 0.5 mM ICG or PA-ICG conjugates were injected intravenously (IV). Blood was drawn retro-orbitally at the specified time points up to 2 days after injection. 2 µL of blood were diluted into 100 µL of PBS and ICG fluorescence was measured using a TECAN Spark 20M microplate reader at excitation and emission wavelengths of 745 and 780-870 nm, respectively. After the 2 day blood draw, mice were euthanized and tumors and organs were excised and imaged with IVIS. The tumor was bisected, half being used for confocal imaging and half for determination of PA content. Tissues were homogenized in gentleMACS™ M Tube containing 1 mL of PBS and equipped with a 600 μm mesh strainer using a gentleMACS dissociator. The resulting tissue homogenates were transferred into a 96-well plate for fluorescence measurement using a microplate reader (TECAN Spark 20M). The percentage of the injected dose was then determined in tumor, spleen, kidney, lung, liver and colon tissues from calibration curves. To prepare these curves, organ tissues from mice without probe injection were homogenized. ICG-conjugated C16 was spiked into tissue samples at different concentrations and the resulting fluorescence intensities were measured using a microplate reader (TECAN Spark 20M).

### Confocal imaging of tumor sections

For Confocal imaging of tumor sections, the tumor was was flash frozen and stored at −80 °C in O.C.T. embedding medium (Sakura Finetek USA Inc, 4583). Tissue was then sectioned with a cryostat (Leica CM 3050s) at 15 μm thickness, then fixed in 4% formaldehyde for 5 min. Tissue sections were counterstained with Hoechst stain (1:4000) for 5 min and imaged on a Nikon CrestOptics X-Light V3 Spinning Disk Confocal microscope with 60x objective using NIS-Elements software. Hoechst staining was imaged at 405 nm, 25% laser power, 100 ms exposure. ICG was imaged at 748 nm, 35% laser power, 1000 ms exposure. mCherry was imaged at 545 nm, 35% laser power, 350 ms exposure.

## Acknowledgements

This project was supported by funding from the Cancer Early Detection Advanced Research (CEDAR) center at the Oregon Health & Science University’s Knight Cancer Institute and the NIH (R21NS135452). GraphPad Prism was used to plot graphs. ChemSketch was used to draw molecular structures. Biorender was used to prepare Figures and TOC.

## References

1. J. Shi, P. W. Kantoff, R. Wooster, O. C. Farokhzad, “Cancer Nanomedicine: Progress, Challenges and Opportunities,” Nat. Rev. Cancer 2017, 17, 20–37.

2. Y. Min, J. M. Caster, M. J. Eblan, A. J. Wang, “Clinical Translation of Nanomedicine,” Chem. Rev. 2015, 115, 11147–11190.

3. S. N. Bhatia, X. Chen, M. A. Dobrovolskaia, T. Lammers, “Cancer nanomedicine,” Nat Rev Cancer 2022, 22, 550–556.

4. P. Zhang, Y. Xiao, X. Sun, X. Lin, S. Koo, A. V. Yaremenko, D. Qin, N. Kong, O. C. Farokhzad, W. Tao, “Cancer Nanomedicine toward Clinical Translation: Obstacles, Opportunities, and Future Prospects,” Med 2023, 4, 147–167.

5. F. Rodríguez, P. Caruana, N. De La Fuente, P. Español, M. Gámez, J. Balart, E. Llurba, R. Rovira, R. Ruiz, C. Martín-Lorente, J. L. Corchero, M. V. Céspedes, “Nano-Based Approved Pharmaceuticals for Cancer Treatment: Present and Future Challenges,” Biomolecules 2022, 12, 784.

6. D. Fan, Y. Cao, M. Cao, Y. Wang, Y. Cao, T. Gong, “Nanomedicine in Cancer Therapy,” Sig. Transduct. Target. Ther. 2023, 8, 293.

7. V. Agrahari, P. Hiremath, “Challenges Associated and Approaches for Successful Translation of Nanomedicines into Commercial Products,” Nanomedicine 2017, 12, 819–823.

8. W. Zhang, D. S. Kohane, “Keeping Nanomedicine on Target,” Nano Lett. 2021, 21, 3–5.

9. J. M. Metselaar, T. Lammers, “Challenges in Nanomedicine Clinical Translation,” Drug Deliv. and Transl. Res. 2020, 10, 721–725.

10. R. Ganpisetti, S. Giridharan, G. S. S. J. Vaskuri, N. Narang, P. Basim, M. R. Dokmeci, M. Ermis, S. Rojekar, A. D. Gholap, N. Kommineni, “Biological Nanocarriers in Cancer Therapy: Cutting Edge Innovations in Precision Drug Delivery,” Biomolecules 2025, 15, 802.

11. H. Cho, S. I. Jeon, C.-H. Ahn, M. K. Shim, K. Kim, “Emerging Albumin-Binding Anticancer Drugs for Tumor-Targeted Drug Delivery: Current Understandings and Clinical Translation,” Pharmaceutics 2022, 14, 728.

12. S. Busatto, S. A. Walker, W. Grayson, A. Pham, M. Tian, N. Nesto, J. Barklund, J. Wolfram, “Lipoprotein-Based Drug Delivery,” Adv. Drug Deliv. Rev. 2020, 159, 377–390.

13. W. Zhang, R. Taheri-Ledari, F. Ganjali, F. H. Afruzi, Z. Hajizadeh, M. Saeidirad, F. S. Qazi, A. Kashtiaray, S. S. Sehat, M. R. Hamblin, A. Maleki, “Nanoscale Bioconjugates: A Review of the Structural Attributes of Drug-Loaded Nanocarrier Conjugates for Selective Cancer Therapy,” Heliyon 2022, 8, e09577.

14. N. N. Pavlova, C. B. Thompson, “The Emerging Hallmarks of Cancer Metabolism,” Cell Metabolism 2016, 23, 27–47.

15. Y. Fu, T. Zou, X. Shen, P. J. Nelson, J. Li, C. Wu, J. Yang, Y. Zheng, C. Bruns, Y. Zhao, L. Qin, Q. Dong, “Lipid Metabolism in Cancer Progression and Therapeutic Strategies,” MedComm 2021, 2, 27–59.

16. E. Frei, “Albumin Binding Ligands and Albumin Conjugate Uptake by Cancer Cells,” Diabetol. Metab. Syndr. 2011, 3, 11.

17. Z. Wang, X. Wang, W. Xu, Y. Li, R. Lai, X. Qiu, X. Chen, Z. Chen, B. Mi, M. Wu, J. Wang, “Translational Challenges and Prospective Solutions in the Implementation of Biomimetic Delivery Systems,” Pharmaceutics 2023, 15, 2623.

18. C. S. Thaxton, J. S. Rink, P. C. Naha, D. P. Cormode, “Lipoproteins and Lipoprotein Mimetics for Imaging and Drug Delivery,” Adv. Drug Deliv. Rev. 2016, 106, 116–131.

19. R. Paliwal, R. J. Babu, S. Palakurthi, “Nanomedicine Scale-up Technologies: Feasibilities and Challenges,” AAPS PharmSciTech 2014, 15, 1527–1534.

20. E. N. Hoogenboezem, C. L. Duvall, “Harnessing Albumin as a Carrier for Cancer Therapies,” Adv. Drug Deliv. Rev. 2018, 130, 73–89.

21. K. K. Ng, J. F. Lovell, G. Zheng, “Lipoprotein-Inspired Nanoparticles for Cancer Theranostics,” Acc. Chem. Res. 2011, 44, 1105–1113.

22. Y. Wang, S.-K. Sun, Y. Liu, Z. Zhang, “Advanced Hitchhiking Nanomaterials for Biomedical Applications,” Theranostics 2023, 13, 4781–4801.

23. B. R. Kimmel, K. Arora, N. C. Chada, V. Bharti, A. J. Kwiatkowski, J. E. Finkelstein, A. Hanna, E. N. Arner, T. L. Sheehy, L. E. Pastora, J. Yang, H. M. Pagendarm, P. T. Stone, E. Hargrove-Wiley, B. C. Taylor, L. A. Hubert, B. M. Fingleton, K. N. Gibson-Corley, J. C. May, J. A. McLean, J. C. Rathmell, A. Richmond, W. K. Rathmell, J. M. Balko, J. T. Wilson, “Potentiating Cancer Immunotherapies with Modular Albumin-Hitchhiking Nanobody– STING Agonist Conjugates,” Nat. Biomed. Eng 2025.

24. S. P. Chawla, Z. Papai, G. Mukhametshina, K. Sankhala, L. Vasylyev, A. Fedenko, K. Khamly, K. Ganjoo, R. Nagarkar, S. Wieland, D. J. Levitt, “First-Line Aldoxorubicin vs Doxorubicin in Metastatic or Locally Advanced Unresectable Soft-Tissue Sarcoma: A Phase 2b Randomized Clinical Trial,” JAMA Oncol. 2015, 1, 1272.

25. P. Famta, S. Shah, N. Jain, D. A. Srinivasarao, A. Murthy, T. Ahmed, G. Vambhurkar, S. Shahrukh, S. B. Singh, S. Srivastava, “Albumin-Hitchhiking: Fostering the Pharmacokinetics and Anticancer Therapeutics,” J. Control. Release 2023, 353, 166–185.

26. L. Xiang, M. R. Stewart, K. Mooney, M. Dai, S. Drennan, S. Holland, A. Quentel, S. Sabuncu, B. R. Kingston, I. Dengos, K. Bonic, F. Goncalves, X. Yi, S. Ranganathan, B. P. Branchaud, L. L. Muldoon, R. F. Barajas, J. M. Fischer, A. Yildirim, “Peptide Amphiphiles Hitchhike on Endogenous Biomolecules for Enhanced Cancer Imaging and Therapy,” Adv. Mater. 2025 Accepted, DOI: 10.1002/adma.202509359.

27. H. Cheng, M. Wang, J. Su, Y. Li, J. Long, J. Chu, X. Wan, Y. Cao, Q. Li, “Lipid Metabolism and Cancer,” Life 2022, 12, 784.

28. C.-F. Deng, N. Zhu, T.-J. Zhao, H.-F. Li, J. Gu, D.-F. Liao, L. Qin, “Involvement of LDL and Ox-LDL in Cancer Development and Its Therapeutical Potential,” Front. Oncol. 2022, 12, 803473.

29. Z.-C. Mo, K. Ren, X. Liu, Z.-L. Tang, G.-H. Yi, “A High-Density Lipoprotein-Mediated Drug Delivery System,” Adv. Drug Deliv. Rev. 2016, 106, 132–147.

30. M. Xiao, J. Xu, W. Wang, B. Zhang, J. Liu, J. Li, H. Xu, Y. Zhao, X. Yu, S. Shi, “Functional Significance of Cholesterol Metabolism in Cancer: From Threat to Treatment,” Exp. Mol. Med. 2023, 55, 1982–1995.

31. A. Sánchez-Iglesias, M. Grzelczak, T. Altantzis, B. Goris, J. Pérez-Juste, S. Bals, G. Van Tendeloo, S. H. Donaldson, B. F. Chmelka, J. N. Israelachvili, L. M. Liz-Marzán, “Hydrophobic Interactions Modulate Self-Assembly of Nanoparticles,” ACS Nano 2012, 6, 11059–11065.

32. A. Dehsorkhi, V. Castelletto, I. W. Hamley, “Self□assembling Amphiphilic Peptides,” J. Pept. Sci. 2014, 20, 453–467.

33. A. Spada, J. Emami, J. A. Tuszynski, A. Lavasanifar, “The Uniqueness of Albumin as a Carrier in Nanodrug Delivery,” Mol. Pharmaceutics 2021, 18, 1862–1894.

34. M. R. Sobansky, D. S. Hage, “Analysis of Drug Interactions with Lipoproteins by High-Performance Affinity Chromatography,” Adv. Med. Biol. 2012, 53, 199–216.

35. A. Hackethal, M. Hirschburger, S. Eicker, T. Mücke, C. Lindner, O. Buchweitz, “Role of Indocyanine Green in Fluorescence Imaging with Near-Infrared Light to Identify Sentinel Lymph Nodes, Lymphatic Vessels and Pathways Prior to Surgery – A Critical Evaluation of Options,” Geburtshilfe Frauenheilkd 2018, 78, 54–62.

36. T. F. Daniels, K. M. Killinger, J. J. Michal, R. W. Wright Jr., Z. Jiang, “Lipoproteins, Cholesterol Homeostasis and Cardiac Health,” Int. J. Biol. Sci. 2009, 474–488.

37. R. Kuai, D. Li, Y. E. Chen, J. J. Moon, A. Schwendeman, “High-Density Lipoproteins: Nature’s Multifunctional Nanoparticles,” ACS Nano 2016, 10, 3015–3041.

38. D. Sleep, J. Cameron, L. R. Evans, “Albumin as a Versatile Platform for Drug Half-Life Extension,” Biochim. Biophys. Acta - Gen. Subj. 2013, 1830, 5526–5534.

39. M. L. James, S. S. Gambhir, “A Molecular Imaging Primer: Modalities, Imaging Agents, and Applications,” Physiol. Rev. 2012, 92, 897–965.

