## Supporting Information for "Lipid Tail Length Determines Nano-Bio Interactions of Peptide Amphiphile Nanostructures"

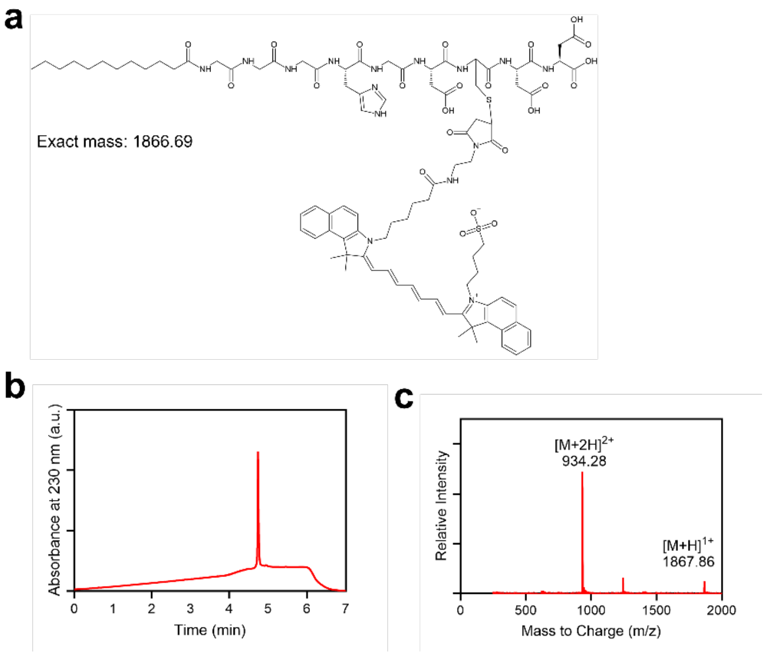


**Figure S1.** Synthesis of ICG-conjugated C12. a) Molecular structure of ICG-conjugated C12. b) LC and c) MS traces of the HPLC purified product.

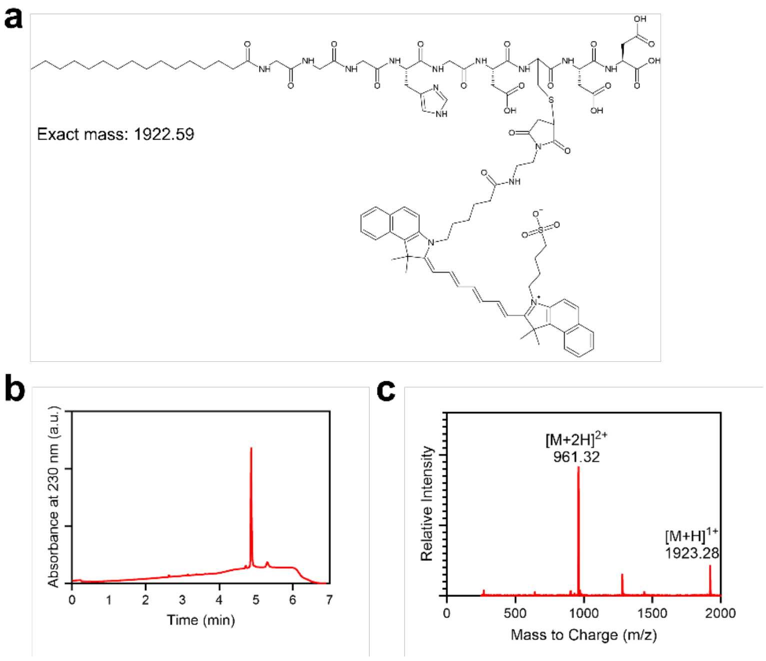


**Figure S2.** Synthesis of ICG-conjugated C16. a) Molecular structure of ICG-conjugated C16. b) LC and c) MS traces of the HPLC purified product.

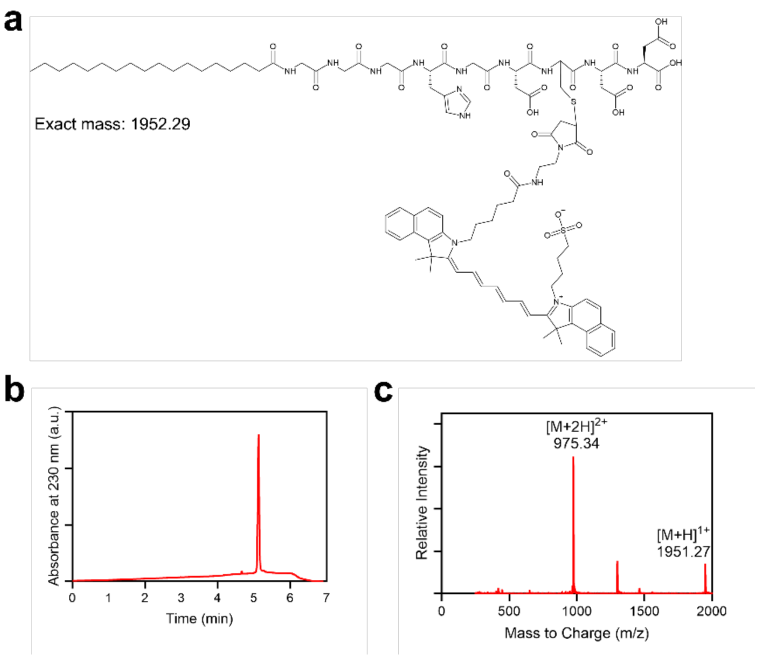


**Figure S3.** Synthesis of ICG-conjugated C18. a) Molecular structure of ICG-conjugated C18. b) LC and c) MS traces of the HPLC purified product.

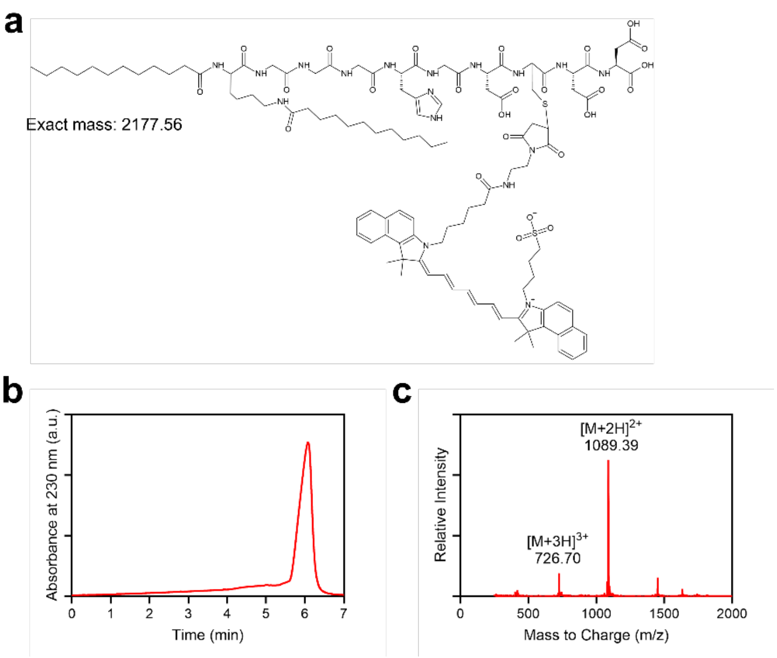


**Figure S4.** Synthesis of ICG-conjugated 2C12. a) Molecular structure of ICG-conjugated 2C12. b) LC and c) MS traces of the HPLC purified product.

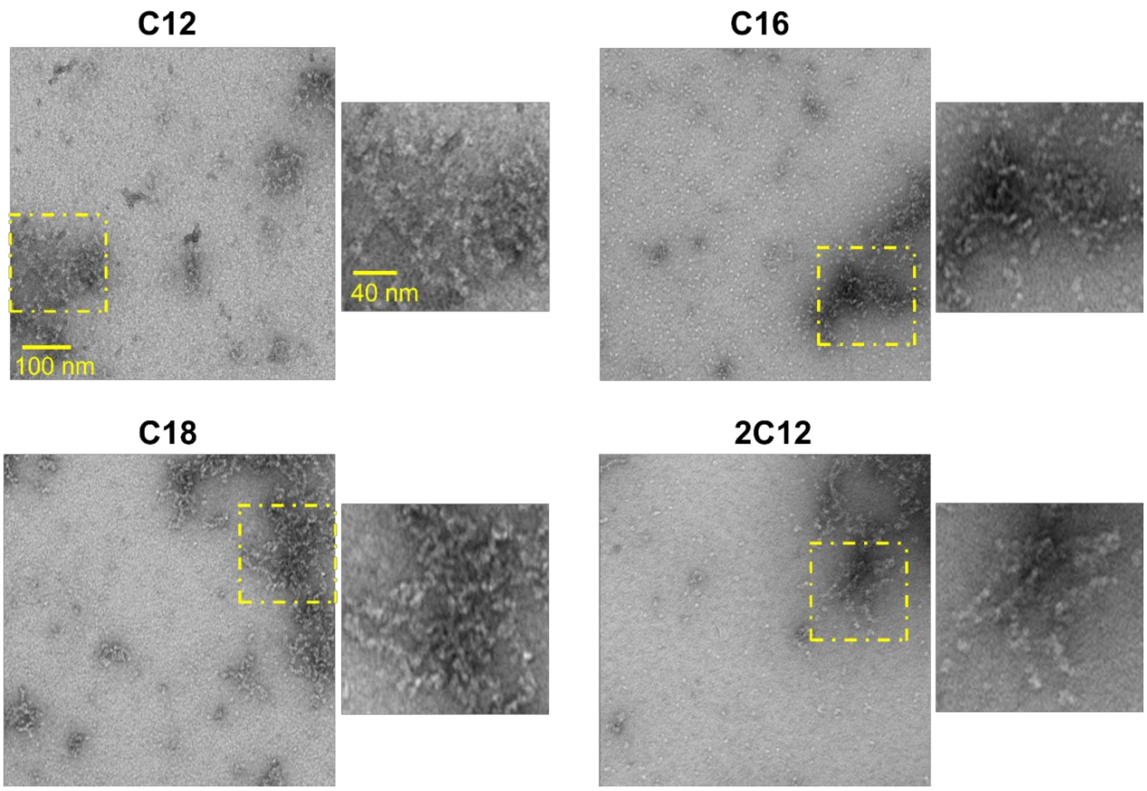


**Figure S5.** TEM images of PAs show that all PAs form micelles in aqueous solution.

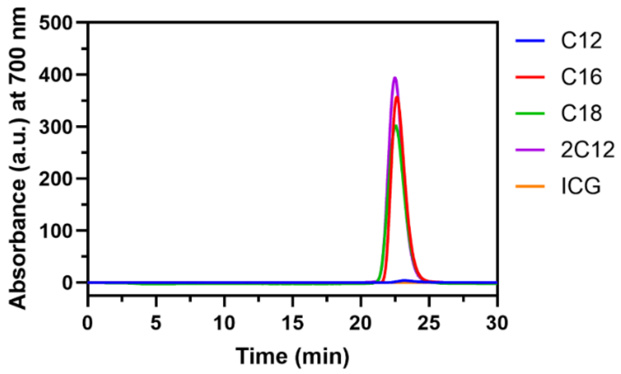


**Figure S6.** FPLC of PAs in PBS. PAs eluted at a single time point without plasma except for C12.

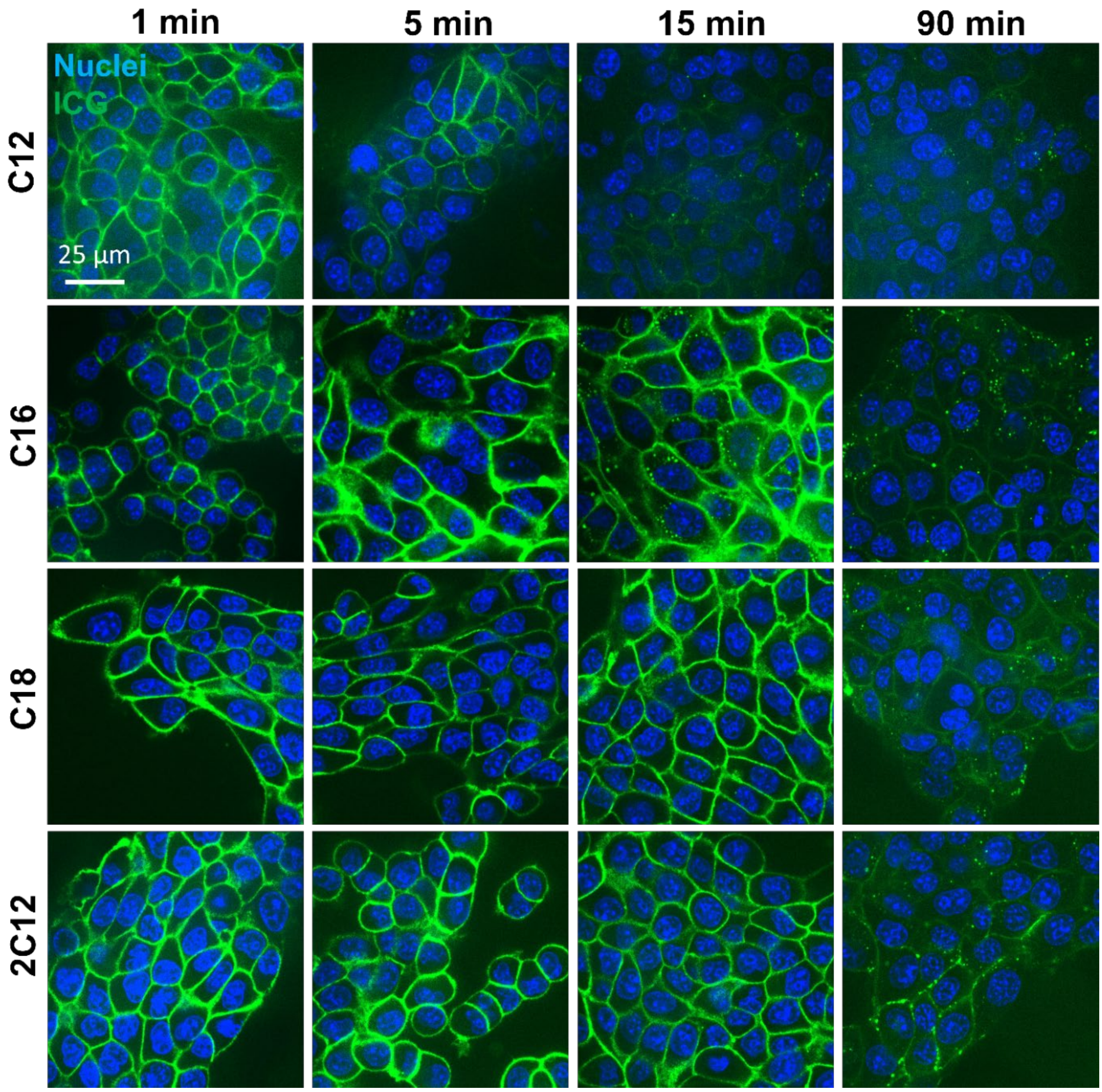


**Figure S7.** Confocal microscope images of 4T1 cells over time show more hydrophobic PAs accumulate in the cell membrane and internalize after longer periods of time than less hydrophobic PAs. Cells were treated with 100 µL of 50 µM PAs for 1 min. Green = ICG, Blue = Nuclei.

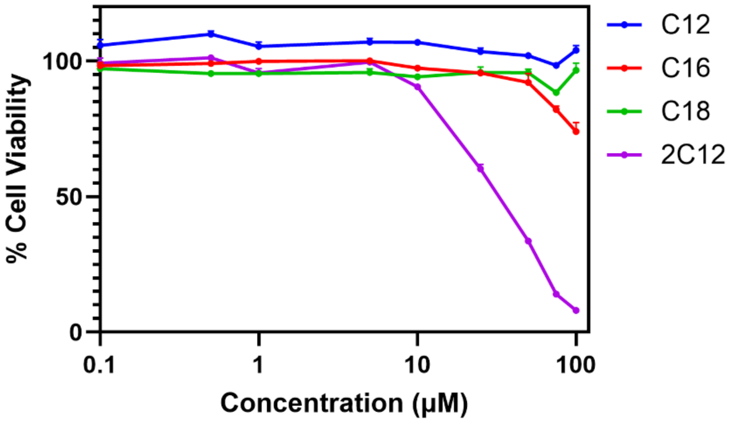


**Figure S8.** Cell viability of 4T1 cells treated with PAs. PAs have little to no effect on cell viability except for 2C12 at concentrations >10 μM. Data are presented as mean ± standard error of the mean (SEM).

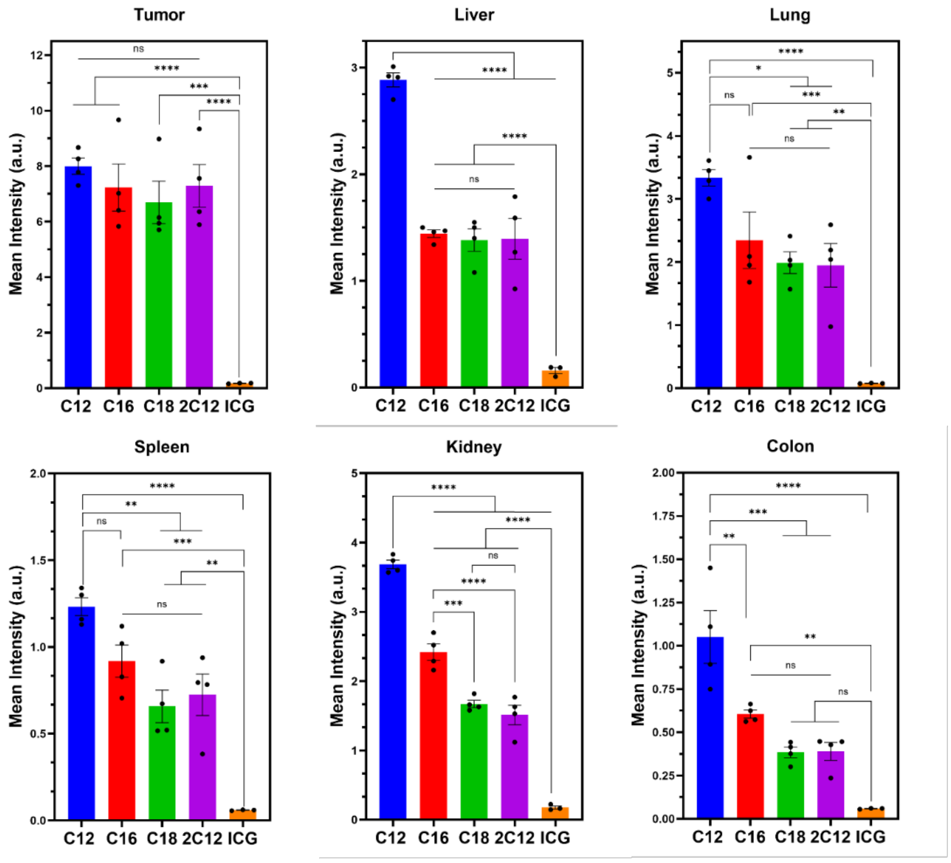


**Figure S9.** IVIS signal of tumors and major organs shows significantly higher tumor accumulation for all PAs compared to free ICG. Error bars are standard error of the mean (SEM). Statistical analysis was performed using one-way analysis of variance (ANOVA) in (b, c). *n.s.* is non-significant and **p* <0.05, ***p* <0.01, ****p* <0.001, *****p* <0.0001.

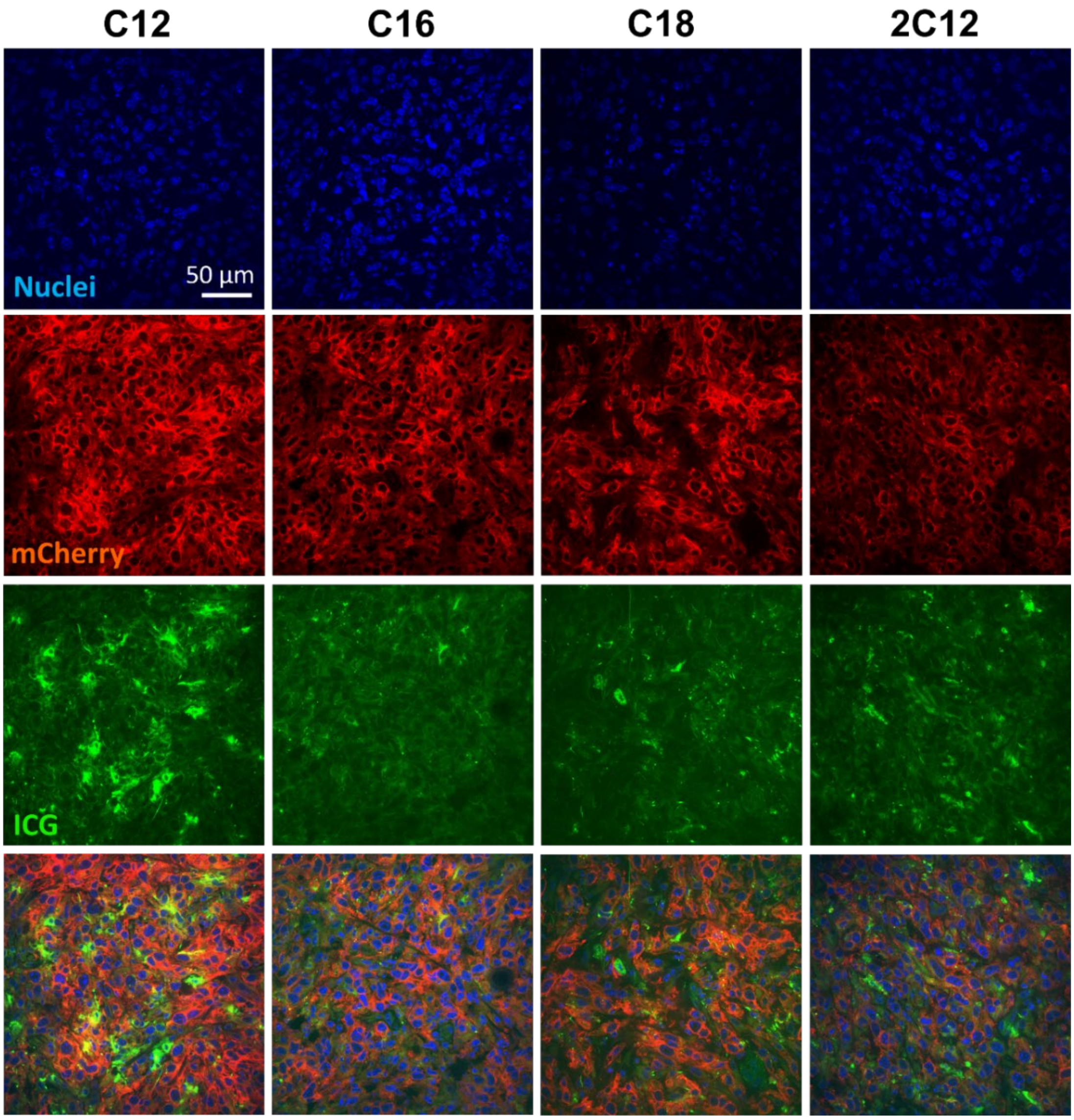


**Figure S10.** ICG-PAs accumulate in cancer cells *in vivo*. Confocal microscopy images of 4T1 mCherry tumor sections from mice treated with ICG-PAs. Tumors were excised from mice 2 days after intravenous injection of 50 nm of ICG-PAs. Green = ICG, Blue = Nuclei, Red = mCherry.

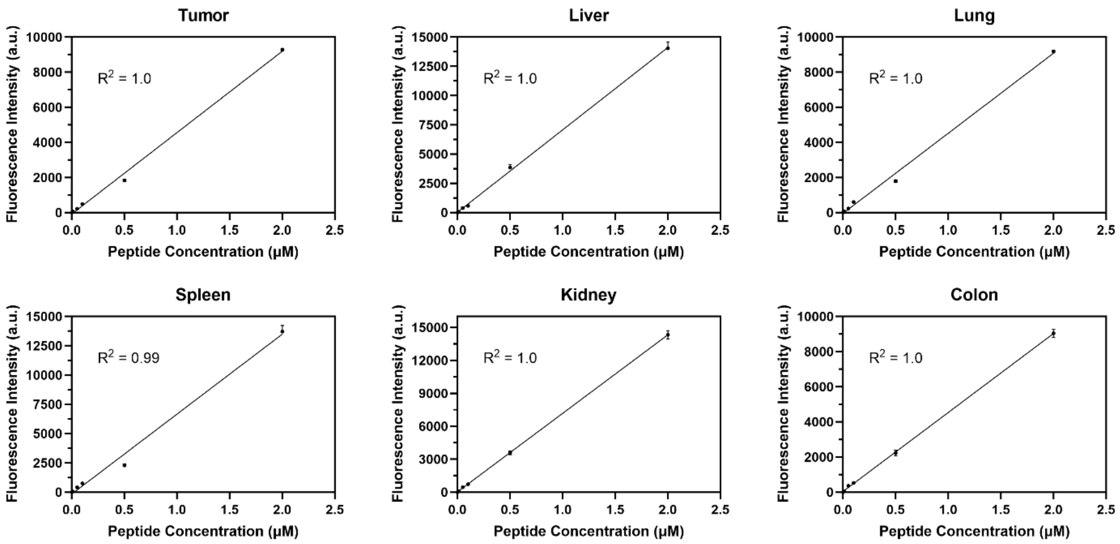


**Figure S11.** Calibration curves of C16-ICG in homogenates of different tissues. Data are presented as mean ± standard error of the mean (SEM).

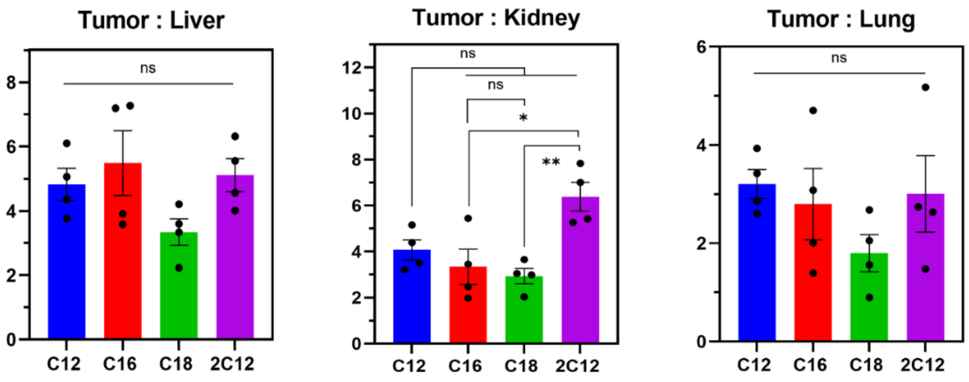


**Figure S12.** Ratio of % ID / gram in tumors versus liver, kidney or lung. Data are presented as mean ± standard error of the mean (SEM). Statistical analysis was performed using one-way analysis of variance (ANOVA). *n.s.* is non-significant and *$p$ <0.05, **$p$ <0.01.
